# Evolutionary Dynamics Do Not Motivate a Single-Mutant Theory of Human Language

**DOI:** 10.1101/517029

**Authors:** Bart de Boer, Bill Thompson, Andrea Ravignani, Cedric Boeckx

## Abstract

**Abstract:** One of the most controversial hypotheses in cognitive science is the Chomskyan evolutionary conjecture that language arose instantaneously in our species as the result of a single staggeringly fortuitous mutation. Here we analyze the evolutionary dynamics implied by this hypothesis, which has never been formalized. The theory supposes the emergence and fixation of a single mutant (capable of the syntactic operation *Merge*) during a narrow historical window as a result of frequency-independent selection under a huge fitness advantage in a population of an effective size that is standardly assumed to have been no larger than ~15 000 early humans. We examine this proposal by combining diffusion analysis and extreme value theory to derive a probabilistic formulation of its dynamics. Perhaps counter-intuitively, a macro-mutation is much more unlikely *a priori* than multiple mutations with smaller fitness effects, yet both hypotheses predict fixation with high conditional probability. The consequences of this asymmetry have not been accounted for previously. Our results diffuse any suggestion that evolutionary reasoning provides an independent rationale for the controversial single-mutant theory of language.

**Significance statement:** In recent years, Chomsky and colleagues have sought support for their minimalist theory of the language faculty from evolutionary considerations. They have argued for a spontaneous emergence of a mutation conferring an advantage for thought independent of communication. Here for the first time a formalization of this view is offered, and contrasted with a more gradual evolutionary scenario. The outcome of our analysis argues against the Chomskyan view.

## 1 Introduction: The Chomskyan Evolutionary Conjecture

The human capacity for language is central to our species’ unique form of intelligence. Understanding the emergence of this capacity is a core challenge for the cognitive and biological sciences. One of the most influential theories of the human language faculty, articulated over decades by Chomsky and associates (1, 2), proposes that Humans are genetically-equipped with a unique computational capacity that specifically allows us to implement computations over hierarchically structured symbolic representations. According to the more recent formulation of this theory, this capacity underpins a syntactic operation known as *Merge,* which is the basis of our ability to represent complex grammars in a way that other species cannot (Chomsky 1995). The strongest version of this theory suggests that the biological foundation of Merge is a single genetic mutation (3–8). Given its staggering consequences, a mutation of this kind is considered a *macro-mutation.* This controversial theory leads to the conclusion that our modern language capacity emerged instantaneously in a single hominin individual who is an ancestor of all Humans. The conclusion follows from a theory-internal hypothesis that *Merge* is either present in full or totally absent (Berwick and Chomsky 2016).

The most detailed articulation of the evolutionary conjecture we will examine here has been presented in Berwick and Chomsky (7). These authors argue that evolutionary considerations represent an independent motivation for the single-mutant theory, because the narrow historical time-window in which the human language faculty must have emerged rules out the emergence and fixation of more than one language-relevant genetic anomaly. This evolutionary motivation for Merge is important because it represents a theory-external rationale for the hypothesis. In this paper, we examine this evolutionary proposal formally by conducting a probabilistic analysis of the evolutionary dynamics that result from its assumptions. Specifically, we formalize the hypothesis that fixation of multiple interacting mutations is less probable than fixation of a macro mutation in this time window, and show that it is wrong.

The key details of the evolutionary conjecture, presented concisely in Berwick and Chomsky (7), are as follows: 1) “universal grammar” (a hypothesized innate basis for the computational capacities implied by their theory of language) can be pared down to little more than a basic combinatorial property known as Merge – this is the major conclusion of the minimalist program (2); 2) only (modern) Humans have Merge; 3) Merge is the result of a single mutation, perhaps provoking a slight rewiring in the brain yielding novel neural circuitry or network configuration that is missing in other species, and this only arose once in our lineage; 4) because it rests on a single mutation, Merge (as a theory of language) has the additional virtue that its genetic basis could be very recent in evolutionary terms, in the sense that Merge has no external prerequisites and therefore the time window between no-Merge and Merge can be small, and this is not true of alternative theories; 5) finally, Chomsky and associates insist that this mutation be kept separate from any considerations of communication: according to their arguments, language is primarily advantageous as a means for internal thought and this is the only phenotypic consequence relevant to the fitness of a hypothetical single-mutant.

Berwick and Chomsky’s account, to date the most detailed articulation of the evolutionary scenario in the mainstream generative research tradition, has been criticized on a variety of grounds (9, 10). Our article does not engage with these criticisms, which dispute arguments 1) through 5). Instead, for the purposes of our analysis, we grant all of Berwick and Chomsky’s assumptions in full, and confine ourselves to examining the consequent evolutionary dynamics formally. In particular, we examine the dynamics of a single, critical, mutation spreading rapidly through a population in a given time window under frequency-independent selection. It is this claim that allows Berwick and Chomsky to argue for a significant selective advantage for Merge --- one that places language at the center of a ‘great leap forward’ view of the human cognitive revolution endorsed by Diamond (11) and Klein (12).

Our evolutionary analysis is made possible by two insights. First, Berwick and Chomsky’s theory is sufficiently precise to place approximate bounds on ordinarily under-determined variables of an appropriate evolutionary analysis, specifically with respect to population size, time-window, and selection regime: these unknown factors can be approximated by combining Berwick and Chomsky’s theoretical proposals with contemporary genetic and demographic findings.

Second, the appropriate degree of belief in Berwick and Chomsky’s proposal has a natural formulation in Bayesian terms, in the sense that it should factor into two distinct quantities: the *a priori* probability of a mutation conferring fitness effects commensurate with their theory of Merge; and the *likelihood* that this mutation would lead to the scenario we observe today – universal acquisition of language among human populations, or *fixation* in the relevant time window. This interpretation enables an examination via separation of the total probability of an evolutionary scenario into: 1) the probability of mutations arising; and 2) the probability of mutations going to fixation (Figure 1). Both of these quantities are well-studied evolutionary phenomena, amenable to approximate analysis. The *a” priori* probability of a mutation can be expressed using results from extreme value theory (13); the *likelihood* of today’s scenario conditional on the Merge-mutant hypothesis can be understood as approximately quantifiable through analysis of *fixation* probabilities: the probability that a single Merge mutant goes to fixation in human populations. Fixation probabilities can be calculated in finite populations via numerous standard methods; here we use diffusion analysis (14, 15).

**Figure 1:**
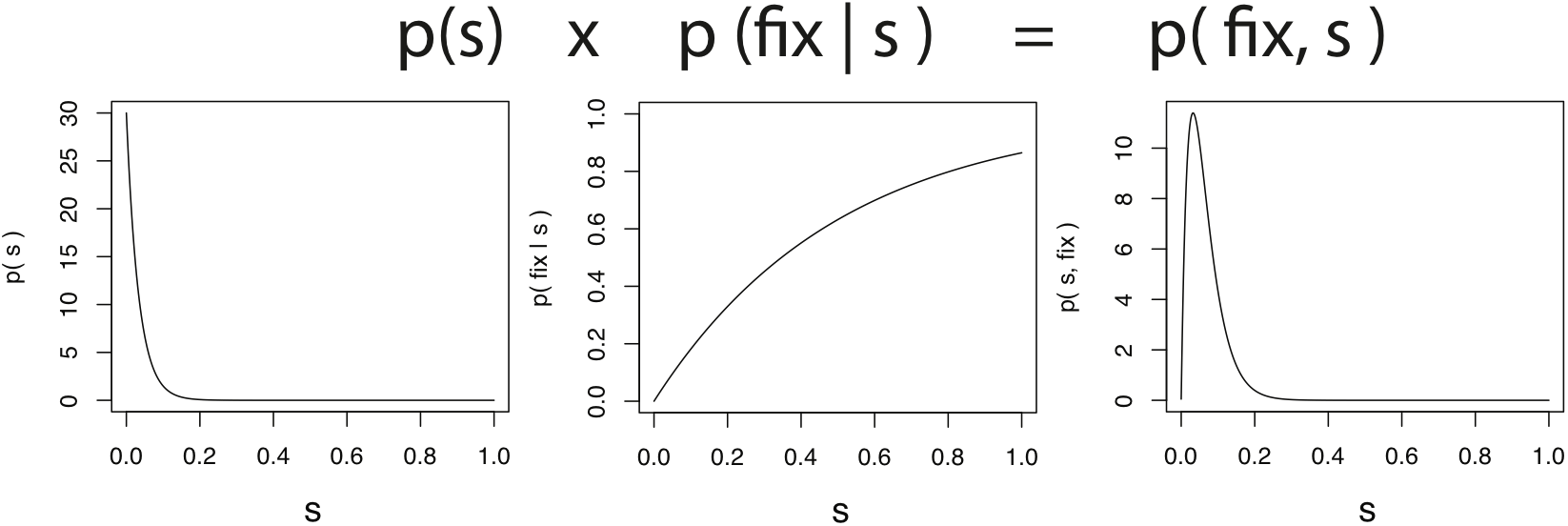
Bayesian perspective on the Berwick-Chomsky conjecture. From left to right: the a priori probability of a mutation of a given fitness advantage occurring, the conditional probability of a given mutation going to fixation (starting with a single mutant) and the total probability of a mutation occurring and going to fixation.

Our analysis contrasts Berwick and Chomsky’s proposal with a scenario in which genetic bases of our linguistic ability evolved through a gradual accumulation of smaller biological changes. We see this as a useful means for model selection among primarily biological theories of the unique basis of language, rather than as a proposal that this kind of account is to be defended per se.

## 2 Model: A Probabilistic Evolutionary Analysis

### 2.1 The probability of one big step versus many smaller steps

According to the single-mutant theory, no other species ever had Merge, and Merge was universally in place before *Homo sapiens* moved out of Africa, restricting its emergence to a time window of approximately 100 000 years. Independent evidence indicates that the effective population size of ancient human populations during this period was maximally around 10 000–15 000 individuals (an estimate of 12800 was published by 16) with considerably smaller bottlenecks (hundreds of individuals). Small effective population sizes are supported by observations of ethno-linguistic group sizes of modern hunter-gatherers, which tend to be around 1500 (17).

Our analysis of Berwick and Chomsky’s proposal examines the fate of a single mutant that arises in a population of this size. Conveniently, fixation of a single mutant in a small finite population is an extensively studied dynamics: theoretical biology provides range of tools for analysis of the core proposal. The single mutation envisaged must have conferred an enormous fitness advantage. Formally, fitness is usually expressed as a *selection coefficient.* Only the ratio of fitness of different individuals is generally relevant, so the wild type (i.e. the individuals without mutations) is defined to have fitness 1. Mutant individuals have fitness 1+s, where s is called the *selection & coefficient*. Because in many cases (as in the case at issue) the organisms under consideration have two copies of each gene, there will be two types of mutant individuals: homozygous (one copy of the gene) and heterozygous (two copies of the gene). These in general have different selection coefficients: *s_aA_* and *s_AA_* respectively in our notation.

Selection is usually assumed to be weak in evolutionary models, which licenses many useful methods of approximate analysis. These methods are unavailable to us as a result of the uncommon assumption that a single mutation confers a large fitness effect. This effect cannot be arbitrarily large, because the number of offspring that an individual could raise in prehistoric and pre-agricultural times was limited by many unrelated factors. One way to balance these constraints is to limit our numerical analyses to selection coefficients s ≤ 1 (which still corresponds to mutants having on average twice the number of offspring than the wild type in the strongest case). Berwick and Chomsky’s assumption that fitness advantages reflect increased capacity for individual thought rather communication imply frequency-independent selection.

Finally, the fitness effect of the proposed mutation would occur in the heterozygote mutant, and its effect must be fully dominant. In other words, both individuals with one copy (heterozygotes) and with two copies (homozygotes) of the mutant gene would have full Merge and thus the full fitness advantage. This is because in the proposed scenario, one either has Merge or not, and therefore there cannot be two separate stages with different fitness effects. The alternative of a recessive mutation, in which individuals need two copies of the mutated gene to have merge, would lead to a strongly reduced probability of the mutation spreading, because homozygotes would be rare initially and the mutant gene would therefore first have to spread through random drift. (Figure 2 provide a sample of probabilities; for an exact description of these calculations see section 2.2). Moreover, we would expect back-mutations in modern populations, which do not seem to occur: Although grammatical deficits are routinely reported in the literature, we are unaware of any claims that there are cases in which Merge has been lost.

**Figure 2:**
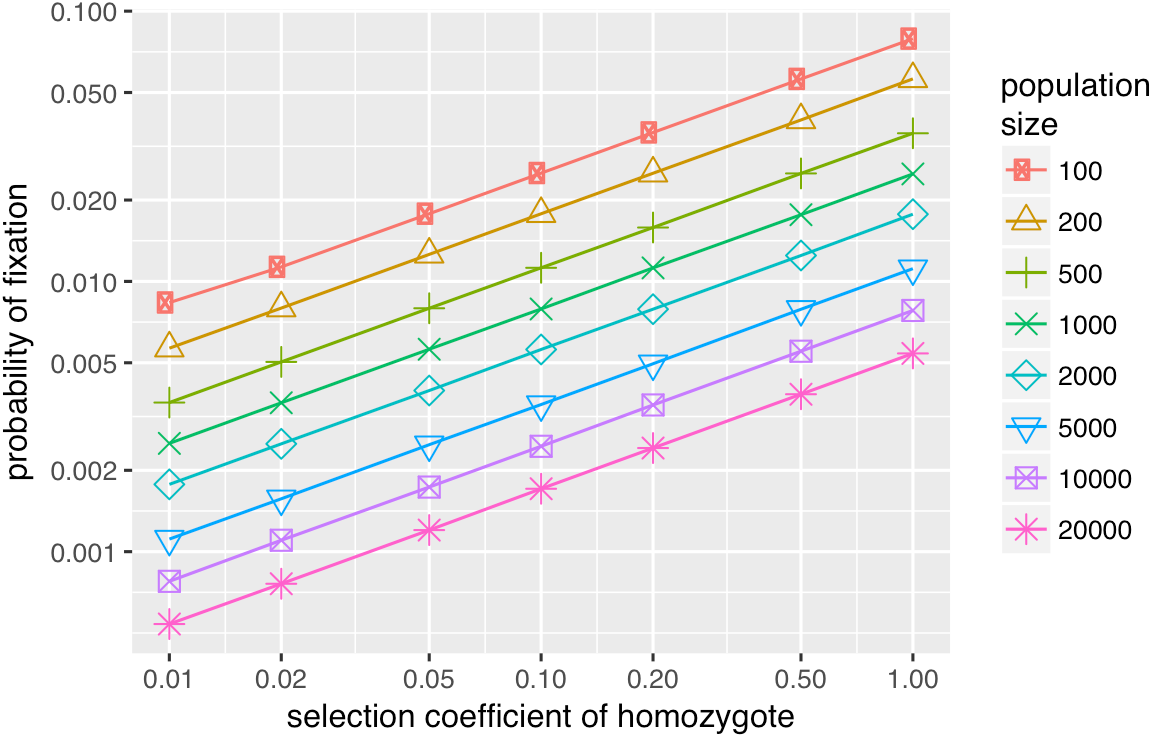
Fixation probabilities for recessive beneficial mutations. Note that these are very low given the large selection coefficients.

By contrast, the alternative scenario for gradual evolution of linguistic ability proposes that the evolution of language happened in a way is far less exceptional by biological standards. This means that there were potentially many mutations involved that had small to moderate fitness effects and that all contributed to the ability for language. Because of the smaller fitness effects, fixation probabilities were lower and fixation times were longer. This does not mean that the overall probability of this scenario is lower, however: multiple mutations evolving in parallel can be present in a population at the same time. If the fitness effects are sufficiently small and additive, mutations can be considered to evolve independently. Also, because the effects of the mutations are small, it is much more probable that if one mutation disappears from the population, eventually another mutation with a similar effect occurs. Finally, in this scenario it is not necessary that all mutations reach full fixation: it is expected that there is some remaining variation in the population. However, the effect of this variation may be small, as there are many genes involved and the variation for a trait determined by many genes with additive effects decreases with the number of genes involved. Unlike the single-mutant hypothesis, there is no reason in this case to assume that the mutations involved were purely dominant. It is unlikely that they were recessive, because recessive mutations of any size are unlikely to spread. However, they may have been semidominant, in which homozygotes have a higher fitness than heterozygotes, which results in higher fixation probabilities and somewhat lower fixation times.

The mutations involved in the gradual scenario may have had positive frequency dependence, because this hypothesis, unlike the Chomskyan view, does not rule out a selective advantage conferred by communication. Positive frequency dependence leads to lower fixation probabilities and longer fixation times, as initially the mutation may not have an influence on fitness. This is quantitatively investigated in section 2.2.4.

### 2.2 Fixation probability and time

Our analysis makes use of the diffusion method developed by Kimura (14, 15). This approach models the proportion of mutants in a population as a continuous value (in reality the proportion is discrete, because the population consists of a finite number of discrete individuals) and it models evolution of the number of mutants as a diffusion process. This approach is flexible enough to model individuals as haploid or diploid, and in the case of diploid individuals it can model different fitness values for heterozygous and homozygous mutants.

#### 2.2.1 General approach

The diffusion approach only considers the evolution of the proportion *p* of mutant alleles *A* (while the wild-type allele is denoted a). Following Kimura (18), a finite population of diploid individuals can be approximated with a diffusion model based on the following backward Kolmogorov equation:

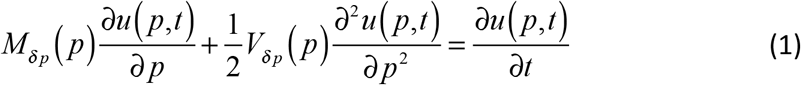

(18, eq. 1), where *u*(*p,t*) is the probability of the mutant allele becoming fixed at time t, if the population starts with a proportion of *p* at time *t* = 0 (where the unit of *t* is generations). *M_δp_*(*p*) is the mean change in proportion *p* per generation, and *V_δp_* (*P*) its variance.

Assuming that there is no more mutation after the initial mutation that generates mutant allele *A*, the mutants will either disappear or take over the population. The probability of this happening corresponds to *u*(*p,∞*) (depending on the boundary conditions, this represents either disappearance or takeover) and as time tends towards large values, the derivative to time of *u* tends to zero. Therefore, in order to know the probability of fixation the following ordinary differential equation needs to be solved:

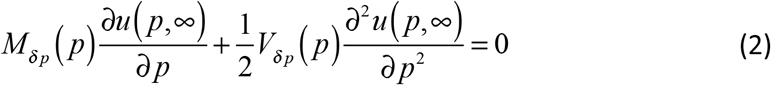

Calculating the probability of the mutant taking over the population requires the boundary conditions:

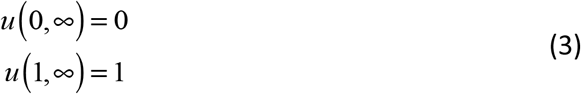

i. e. if the population starts without mutants, the probability of them taking over is zero, and if the population starts with all mutants, they have already taken over, so the probability of takeover is 1.

#### 2.2.2 Mean and Variance Terms

There are different ways of approximating the mean and variance for a given population, and these result in different expressions for *M_δp_*(*p*) and *V_δp_*(*p*). The simplest way is to model the population as if it were haploid (see Supplementary section S1 for a demonstration that this way is also the one that most closely approximates Monte Carlo simulations). The reason that this works even for diploid organisms is that initially, homozygous mutants will be very rare (in a randomly mixing population of sufficient size). Therefore, the mutant allele may become so frequent before homozygotes play a role in its evolution, that it is highly likely that the mutant allele reaches fixation. We can therefore ignore the existence of heterozygotes. The mean change and its variance (see Supplementary section S2 for a derivation) are then:

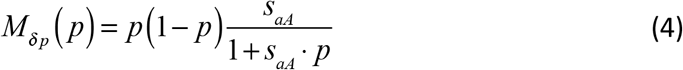

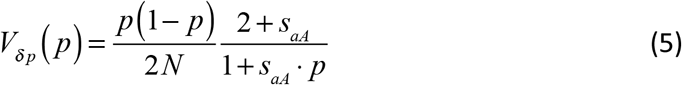

where *s_aA_* is the selection coefficient of the heterozygote mutant (i.e. assuming the fitness of the wild type is 1, the fitness of the heterozygote mutant is 1+*s_aA_*) and *N* is the population size. Assuming a haploid population is mathematically equivalent to assuming a diploid population with *s_AA_* = *2s_aA_*, (this is also called the case of no dominance or semidominance).

This simple approximation also has a closed form solution for the fixation probability:

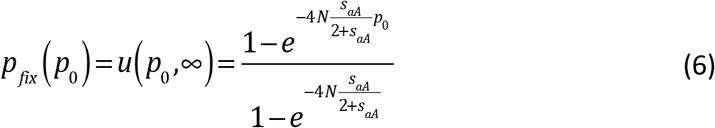

where *p*_0_ is the initial frequency of mutant alleles (which is *1/2N* when starting with a single heterozygous mutant).

#### 2.2.3 Fixation time

In order to estimate the time it takes for a mutant to reach fixation in a population (starting with a given proportion of mutants) the Fokker-Planck equation (2) can be adapted. Setting it equal to minus the fixation probability (instead of 0) results in an equation for the conditional fixation time *θ*(*p*), i.e. the product of the probability of reaching fixation multiplied by the time needed to reach fixation, starting with a proportion *p* of mutants (19, equations XII.3.4a and XII.3.4b):

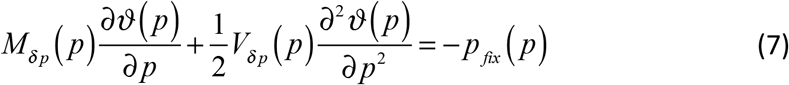

and using boundary conditions:

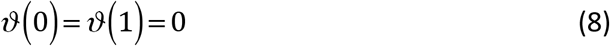

The unconditional fixation time *τ*(*p*) can then be calculated by dividing the conditional fixation time by the fixation probability. Although ignoring the fact that the population consists of heterozygotes may work well for calculating fixation probabilities, it is not expected to work well for calculating fixation times. This is because it is assumed in our analysis that homozygote mutants have no advantage over heterozygote mutants, while approximating the population as haploid implicitly assumes that homozygotes have a selection coefficient that is twice that of heterozygotes. For this reason Kimura’s (18) approximation is used:

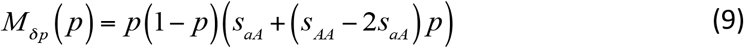

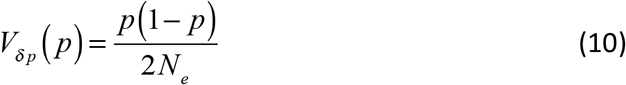

where *N_e_* is the *effective population size,* i.e. the size of a randomly mating population that behaves the same as the actual population (which may not have random mating). These appear to give satisfactory values compared to the Monte-Carlo simulation (see Supplementary section S1).

Solving the resulting equations numerically, we obtain fixation times, expressed in generations (this can be converted to years by multiplying with generation time - the mean time between generations in years - which is about 25–30 years, 20) shown in Figure 3. It shows that for small values of *N·s*, fixation time is approximately 4N, which corresponds to the theoretical value for drift (21). For larger values, fixation time drops rapidly.

**Figure 3:**
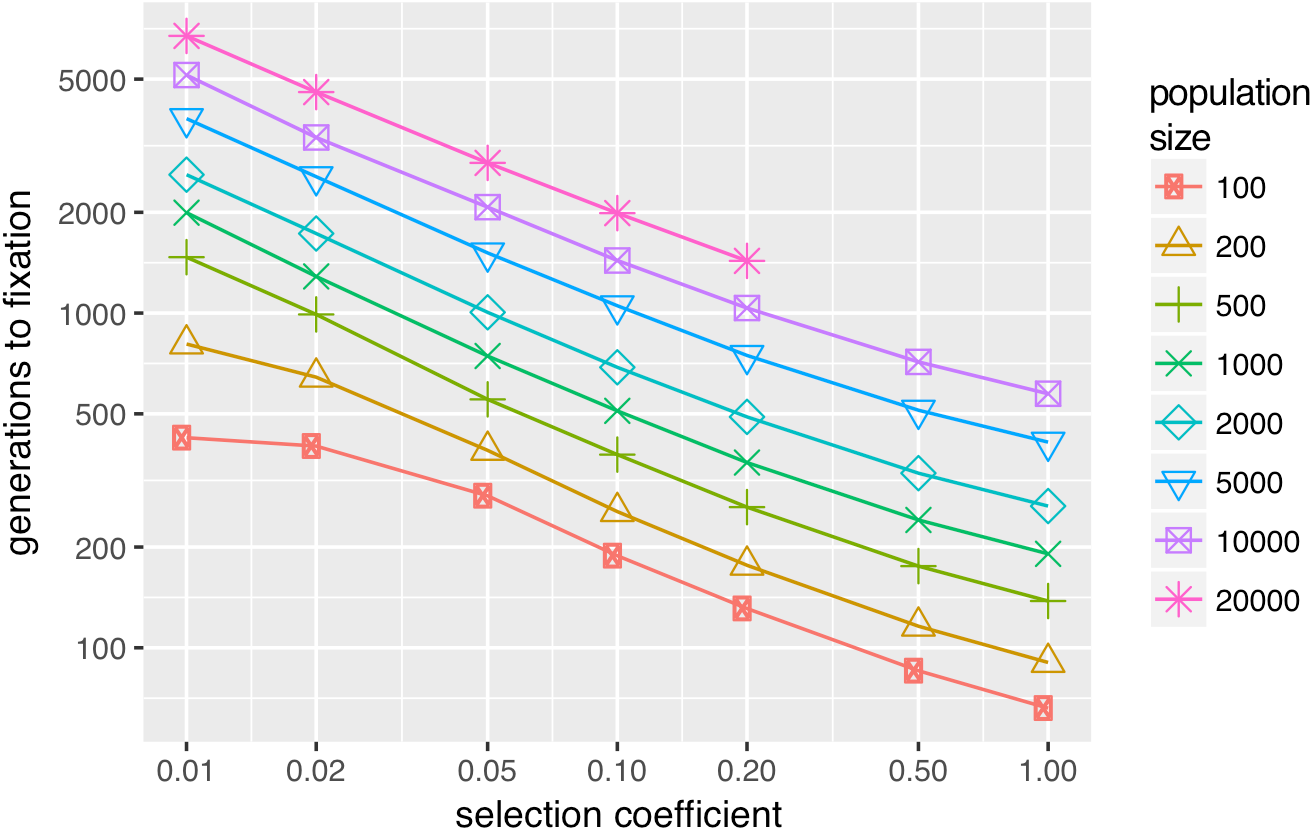
Unconditional mean fixation times for different selection coefficients and population sizes (in individuals, so the number of alleles is twice the number). Note that the missing points in the graph could not be calculated because of numerical limits. Generations can be converted in years by multiplying with generation time, which is from 25–30 years (20).

#### 2.2.4 Frequency dependent fitness

When fitness is (positively) frequency dependent, fixation probabilities are expected to become smaller, as initially the spread of mutants will only be due to drift; if the mutation is rare, individuals that have it will have little or no advantage of it. A simple model for frequency dependent fitness is to make the fitness of a mutant individual linearly dependent on the frequency *p* of mutants in the population. The equations for the selection coefficients then become:

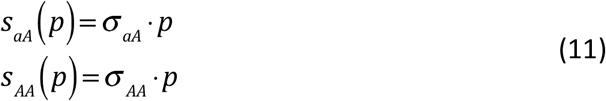

where *s_aA_*(*p*) and *s_AA_*(*p*) are the selection coefficients of the heterozygote respectively the homozygote and *σ_aA_* and *σ_AA_* are the slope of the corresponding selection coefficients. These frequency dependent selection coefficients can be substituted in the equations for the mean change and the variance in (2) and (7), and the fixation probabilities (and fixation times) can be estimated.

As can be seen in Figure 4 (which uses *σ_AA_* = *2σ_aA_* to be as close as possible to the semidominant case) the fixation probabilities become dramatically lower in the case of frequency dependent selection, even for high values of the slope of the selection coefficients (implying high values of the selection coefficients when mutant frequencies become sufficiently high).

**Figure 4:**
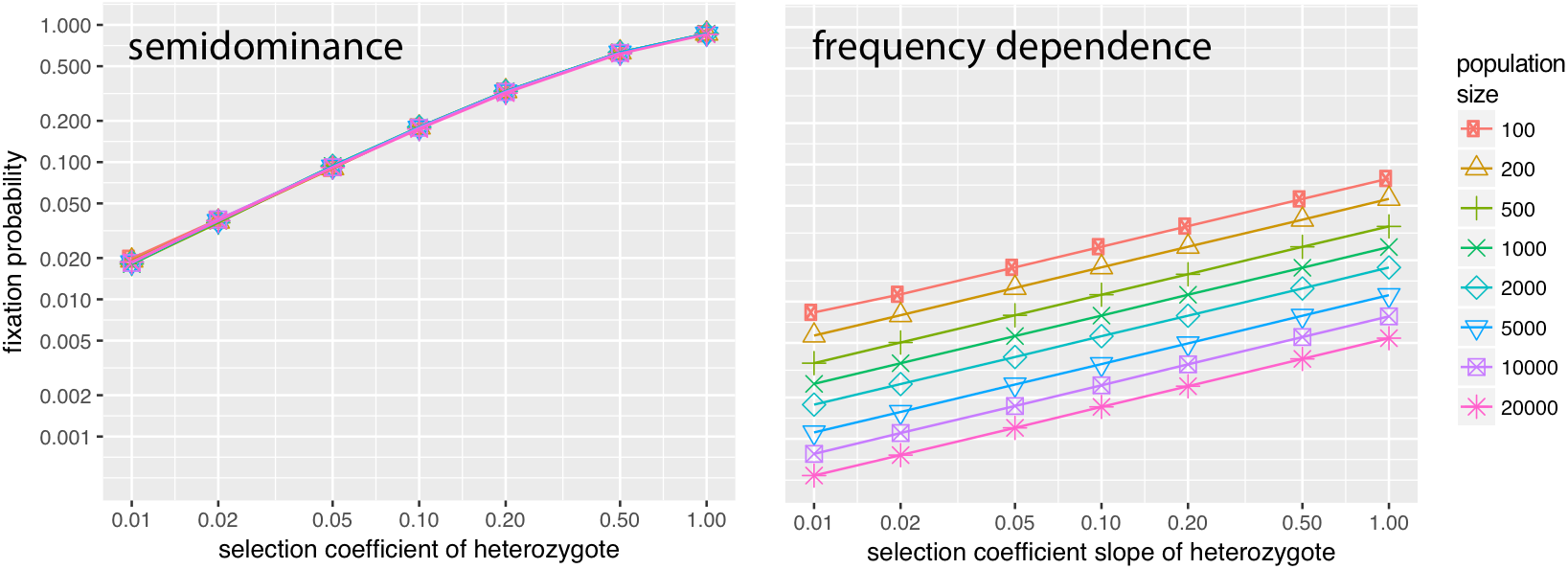
Comparison of the fixation probabilities of the semidominant case and the case where fitness has positive (linear) frequency dependence. Note that the vertical axis (of fixation probabilities) is logarithmic, and that therefore the fixation probabilities in the frequency dependent case are much lower than in the semidominant case.

#### 2.2.5 Time and probability conclusion

Fixation probability can be modeled with satisfactory accuracy (given that our model parameters are approximations anyway) by using the haploid/semidominant approximation. Fixation time does need to take into account whether the mutation under investigation is dominant, recessive or otherwise. However, assuming the proposed mutation for Merge was dominant and conveyed a large fitness advantage, neither fixation probability nor fixation time are obstacles to the theory. The many mutations with small fitness effects proposed by the gradual scenario would take longer to reach fixation and would be less likely to do so. However, as we already remarked above, multiple mutations of this kind could spread at the same time, and importantly, it is less problematic for this scenario if one mutation does not reach fixation, because it is expected that similar mutations will recur, while the proposed macromutation for Merge will not.

To revisit the Bayesian interpretation we laid out earlier, an appropriate summary of the preceding findings is that as far as fixation probabilities go, both hypotheses (single-mutant and gradual evolution) assign a realistically high likelihood to the observation that language is universal in our species (i.e. the mutations have spread to fixation or near fixation), each for different reasons. In other words, consideration of fixation dynamics does not decisively choose between these two competing hypotheses; both are plausible routes to fixation in populations of approximately the hypothesized size and time depth, conditional on the emergence of the relevant mutation(s) in the first place. As a result, evolutionary support for these hypotheses turns exclusively on consideration of their *a priori* probability.

### 2.3 Probability of mutations

Orr offers an appropriate theoretical result for the distribution of the size of beneficial mutations, based on extreme value theory (13). Orr assumes that there are a potentially large number of variants of the genome, all with fitness effects that have been drawn from a distribution that fulfills reasonable assumptions (more specifically, the distribution has to be of the Gumbel type). He further assumes that the wild type already has relatively high fitness compared to all possible variants, because it already is the result of many generations of selection. There are therefore only comparatively few variants that would result in higher fitness. In this case it can be shown (using extreme value theory) that the fitness of beneficial mutants follows an exponential distribution, and that the parameter of this distribution is equal to the reciprocal of the expected value of the fitness difference between the best and second best mutants. This leads to the following expression for the probability *p* of a fitness effect:

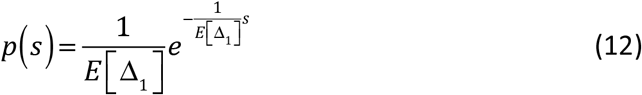

where *s* is the increase in fitness, which happens to be equivalent to the selection coefficient used in section 2.2 and *E*[Δ_1_] the expected value of the difference in fitness between the best and the second best mutations. Experimental evidence is in general agreement with the predictions implied by this model (reviewed by 22) but because advantageous mutations are so rare the overall evidence is weak and could also fit other potential distributions. Eyre-Walker and Keightley also point out a potential weakness in Orr’s analysis: it assumes that fitness values are drawn from a constant distribution, but in reality this distribution may be variable over time. A computational model by Cowperthwaite et al. (23) lends support to this criticism: it shows that in their model there are many more small effect mutations, and only when *“the vast majority”* of these are ignored, does the distribution of fitness effects become exponential. It also shows that the distribution of mutant fitness depends on the parent fitness, lending support to the hypothesis that adaptation to the fitness landscape modifies the distribution of mutant fitness.

There are at least three reasons why in the analysis presented here an exponential distribution is nevertheless reasonable. The first is pointed out by Eyre-Walker and Keightley (22, p. 614): *“Although the Gillespie–Orr prediction* [i.e. the exponential distribution of mutant fitness] *might not be correct for all advantageous mutations, it may apply to mutations of large effect. Such large-effect mutations seem to be those that contribute most to adaptation.”* This is related to the second reason: the single-mutant hypothesis proposes a mutation of very large effect, and the probability of this would lie well within the exponential region. Assuming an exponential distribution and ignoring all beneficial mutations of small effect likely overestimates the probability of the emergence of a ‘Merge mutant’, and is therefore a conservative approximation. The final reason is purely pragmatic: there is no closed form (nor even a clear empirical form) of the true distribution of mutant fitness, and the exponential distribution is relatively easy to work with.

In order to use the exponential distribution to estimate the probability of an evolutionary scenario, it is necessary to have an estimate of the parameter Δ_1_. For this it is necessary to have data about the fitness effects of beneficial mutations, and in particular data about the distribution of such mutations when they occur, as opposed to after they have gone to fixation (which would conflate their probability of occurrence with the probability of going to fixation). For practical reasons, data of this kind are hard to obtain, and only exist for very rapidly evolving organisms such as viruses. A small data set is presented in Sanjuán et al. (24). Fitting an exponential distribution to their datapoints of “random” mutations (which are the ones needed to estimate Δ_1_) a value of *E*[Δ_1_] of between 1/53 (for maximum likelihood estimation) to 1/37 (for least sum of squares estimation) is found. The data, the interpolation and the theoretical prediction are given in Figure 5.

**Figure 5:**
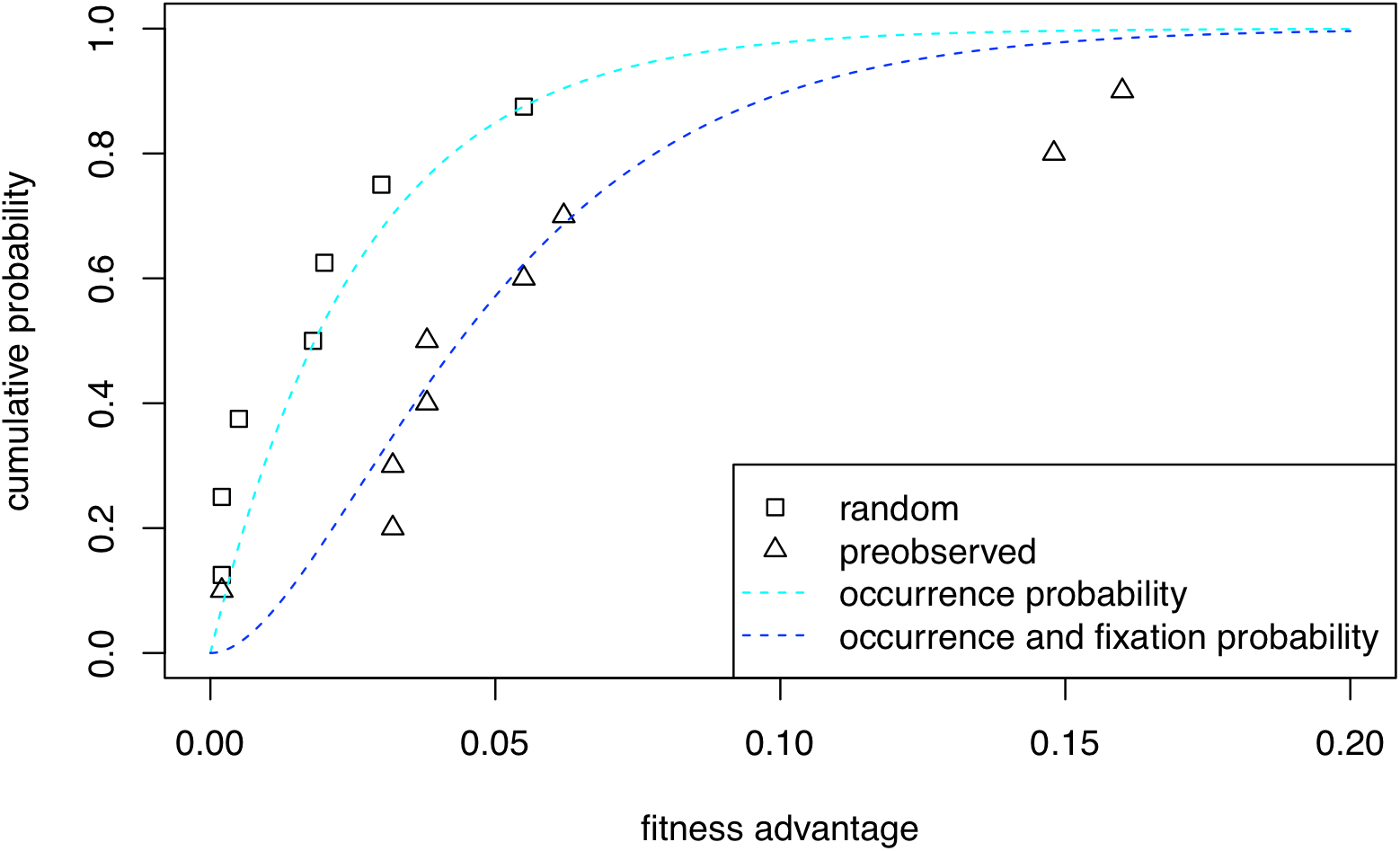
Fitness advantage of viruses from Sanjuán et al.’s (24) sample, interpolated curve (minimal sum of squares fit) on the random data points and theoretically predicted curve of the preobserved (occurrence and going to fixation) probabilities. For these curves a large effective population size was assumed, so the term with N in equation (17) was ignored.

Given the small sample, and given that it is unknown how well the data for viruses map on data for humans, there is considerable uncertainty in this estimate. Because lower values correspond to lower probability of large mutations, and in order to prevent an unjustified bias against large mutations, an estimate of 1/30 for the parameter of the exponential distribution is a reasonable approximation.

### 2.4 Probability of the two scenarios

In order to compare the two scenarios, it is necessary to calculate the probability of a single mutation occurring and going to fixation. For clarity’s sake the term *improvement* is introduced for the event of a beneficial mutation occurring *and* going to fixation. The magnitude of the improvement is defined to be equal to the fitness advantage of the mutation. Finally, the probabilities of single improvements must be combined in order to calculate how many improvements are needed in order to obtain the desired trait.

#### 2.4.1 The probability of a single improvement

The probability of a single improvement is the product of a mutation of a given fitness advantage occurring (equation (12), and that mutation going to fixation. It has been shown that equation (6) gives a satisfactory approximation of the latter probability. This results in the following expression for the probability density function (with appropriate renaming of the variables, *α* = 1/*E* [△_1_] and a scaling factor *β* to guarantee it integrates to 1):

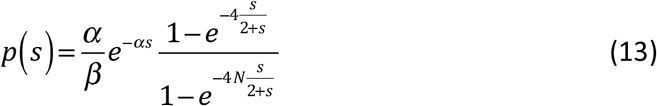

This expression is somewhat unwieldy, as it does not appear that there is a closed form solution to its integral. However, observing that because *α* is large the probability drops off quickly, such that most improvements occur for small *s*, the fixation probability term can be simplified to:

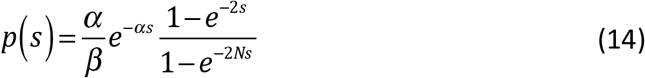

but although this can be integrated, it does not result in a very useful closed form solution. Replacing the second term by 1–*e*^2*s*^ results in a good approximation for *Ns* 0, but unfortunately, it assigns zero probability to mutations with a very small selection coefficient (this should be 1/*N*). However, adding a correction term of *e*^−(*N*+1)*s*^/*N* results in an excellent approximation for all *s*, resulting in our final equation for the probability of an improvement:

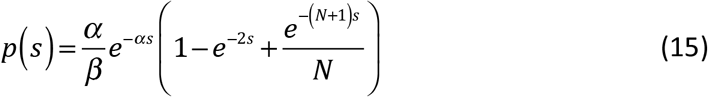

which is straightforward to integrate for calculating the scaling factor *β* and the cumulative distribution function. This results in the following equations. The scaling factor is:

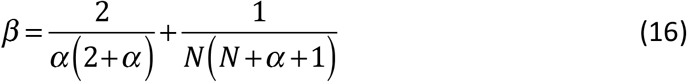

The cumulative distribution function is:

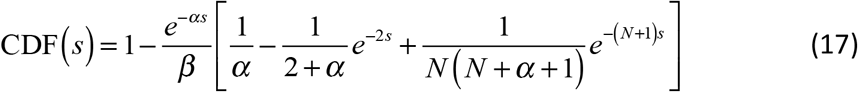

That this equation is a good approximation is illustrated by the fit to the experimental observations illustrated with the dark blue line in Figure 5.

#### 2.4.2 The expected number of improvements necessary

The distributions of improvements can be used to estimate how many improvements are necessary to achieve a given total improvement *m* and what the maximum size of improvements was. The (mean) number of improvements necessary is not equal to *m* divided by the mean size of an improvement, but somewhat larger: because improvements come in discrete steps, the total improvement achieved will in general exceed *m* by a little bit. Therefore it is easier to simulate this with Monte-Carlo techniques. This is implemented by accumulating random improvements until they exceed the threshold *m*. For *m* = 1, *α* = 30 and *N* = 1500, the histograms of the number of improvements and the maximum size of improvements in each string of improvements are given in Figure 6.

**Figure 6:**
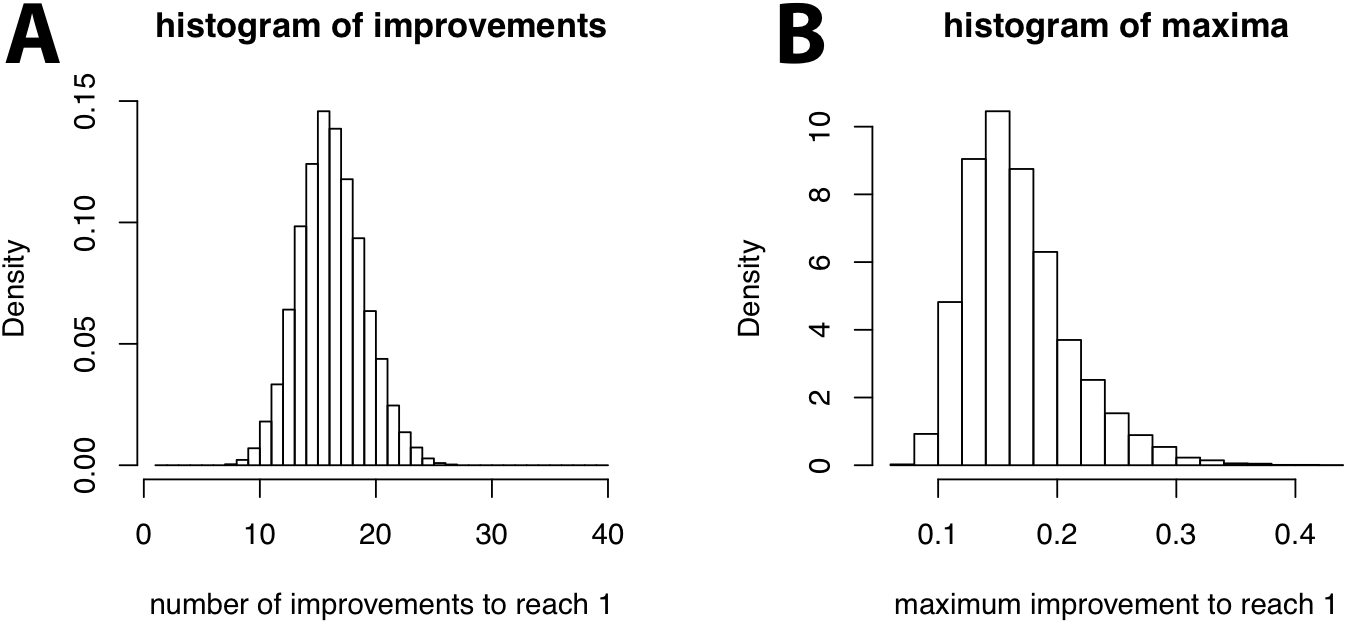
Histograms of the number of improvements needed to reach a total improvement of 1 (A) and the maximal step size that occurred in each run (B), Parameters were α = 30 and N = 1500.

The following observations can be made: first, the number of necessary steps is larger than 1. Even for a modest (total) improvement (e.g. 1), it is exceedingly unlikely that an improvement of this magnitude would occur in a single step. Second, relatively large improvements are common (when relaxing the restriction that they have arisen in one step). For almost all simulations, improvements with large effects emerged. This is in line with the observation by Eyre-Walker and Keightley that “[…] *large-effect mutations seem to be those that contribute most to adaptation.”* (22, p. 614). Even though beneficial mutations of large effect are rare, they are more likely to go to fixation. This is related to our final observation, that the total number of improvements involved is generally not very large. In summary: a single mutation event is ruled out in probabilistic terms. Instead, our analysis favors an evolutionary scenario with relatively few, but relatively important events that unfold gradually (see supplementary material section 3 for the precise relation between the number of mutations required and parameters *m* and *α*).

## 3 Discussion

We have applied a variety of techniques from theoretical biology to the question of how to quantify the probability of a complex trait like language evolving in a single step, in many small steps, or in a limited number of intermediate steps, within a specific time window and population size. The dynamics we studied shows that a limited number of intermediate steps is most probable. On these grounds we conclude that the single-mutant hypothesis is not independently motivated by evolutionary dynamics.

This conclusion challenges a key Chomskyan argument in favor of Merge and the strongest minimalist hypothesis (7), because we have shown that it is not correct to argue that fixation of multiple mutations is less probable than fixation of a single macro-mutation. By diffusing the proposed evolutionary rationale for a single, undecomposable computational innovation, we challenge one of the central theory-external motivations for Merge. Combined with mounting evidence from the archeological record that goes against a very recent ‘great leap forward’ (25, 26) and that indicates that the ability for language is older and evolved over a longer time than hitherto thought (27–29), as well as the rarity of truly fixed mutations in the modern human lineage (30–32), we are inclined to argue that evidence favors the view that gene-culture co-evolution is a more compelling approach to the human language faculty. This view predicts that any innate predispositions for language amount to defeasible inductive biases, each of which weakly constrains behavior on its own, but makes a significant contribution thanks to the intervention of culture (33, 34), and therefore re-frames the evolutionary explanandum to include the social and cognitive conditions that facilitate culture (35, 36).

## Supporting information

Supplementary Sections

## Acknowledgments

**de Boer**

ERC Proof of Concept Grant AI-CU [grant number 789999]

**Thompson**

Max Planck Institute Temporary postdoc grant

**Ravignani**

European Union Horizon 2020 research and innovation programme Marie Skłodowska-Curie grant, agreement No 665501 with the research Foundation Flanders (FWO) (Pegasus2 Marie Curie fellowship 12N5517N)

**Boeckx**

1. Spanish Ministry of Economy [FFI-2106-78034-C2-1-P/FEDER]
2. Catalan Government [2017-SGR-341]
3. Japanese MEXT/JSPS Grant-in-Aid for Scientific Research on Innovative Areas 4903 (Evolinguistics: JP17H06379)

## References

1. Chomsky N (1965) Aspects of the Theory of Syntax (MIT Press, Cambridge, Mass.).

2. Chomsky N (1995) The minimalist program (MIT Press, Cambridge, MA).

3. Berwick RC, Chomsky N (2011) The Biolinguistic Program: The Current State of its Evolution. The Biolinguistic Entreprise: New Perspectives on the Evolution and Nature of the Human Language Faculty, eds Di Sciullo AM, Boeckx C (Oxford University Press, Oxford), pp 19–41.

4. Berwick RC, Friederici AD, Chomsky N, Bolhuis JJ (2013) Evolution, brain, and the nature of language. Trends Cogn Sci 17(2):89–98.

5. Bolhuis J, Tattersall I, Chomsky N, Berwick RC (2014) How Could Language Have Evolved? PLOS Biol 12(8):e1001934.

6. Everaert MB, Huybregts MA, Chomsky N, Berwick RC, Bolhuis JJ (2015) Structures, not strings: linguistics as part of the cognitive sciences. Trends Cogn Sci 19(12):729–743.

7. Berwick RC, Chomsky N (2016) Why only us: Language and evolution (MIT press).

8. Friederici AD, Chomsky N, Berwick RC, Moro A, Bolhuis JJ (2017) Language, mind and brain. Nat Hum Behav 1(10):713.

9. Boeckx C (2017) Not only us. Inference Intern Rev Sci 3(1).

10. Fitch WT (2017) On externalization and cognitive continuity in language evolution. Mind Lang 32(5):597–606.

11. Diamond J (1998) Guns, germs and steel: a short history of everybody for the last 13,000 years (Random House).

12. Klein RG (2009) The human career: human biological and cultural origins (University of Chicago Press, Chicago, Il).

13. Orr HA (2003) The distribution of fitness effects among beneficial mutations. Genetics 163(4):1519–1526.

14. Crow JF, Kimura M (1970) An Introduction to Population Genetics Theory (The Blackburn Press, Caldwell, NJ).

15. Kimura M (1957) Some problems of stochastic processes in genetics. Ann Math Stat 28(4):882–901.

16. Fagundes NJ, et al. (2007) Statistical evaluation of alternative models of human evolution. Proc Natl Acad Sci 104(45):17614–17619.

17. Marlowe FW (2005) Hunter-Gatherers and Human Evolution. Evol Anthropol 14(1):54–67.

18. Kimura M (1980) Average time until fixation of a mutant allele in a finite population under continued mutation pressure: Studies by analytical, numerical, and pseudo-sampling methods. Proc Natl Acad Sci 77(1):522–526.

19. van Kampen NG (1992) Stochastic Processes in Chemistry and Physics (Elsevier Science B. V., Amsterdam).

20. Langergraber KE, et al. (2012) Generation times in wild chimpanzees and gorillas suggest earlier divergence times in great ape and human evolution. Proc Natl Acad Sci 109(39):15716.

21. Kimura M, Ohta T (1969) The Average Number of Generations until Fixation of a Mutant Gene in a Finite Population. Genetics 61(3):763–771.

22. Eyre-Walker A, Keightley PD (2007) The distribution of fitness effects of new mutations. Nat Rev Genet 8(8):610.

23. Cowperthwaite MC, Bull JJ, Meyers LA (2005) Distributions of Beneficial Fitness Effects in RNA. Genetics 170(4):1449.

24. Sanjuán R, Moya A, Elena SF (2004) The distribution of fitness effects caused by single-nucleotide substitutions in an RNA virus. Proc Natl Acad Sci 101(22):8396–8401.

25. McBrearty S, Brooks AS (2000) The revolution that wasn’t: a new interpretation of the origin of modern human behavior. J Hum Evol 39(5):453–563.

26. Scerri EML, et al. (2018) Did Our Species Evolve in Subdivided Populations across Africa, and Why Does It Matter? Trends Ecol Evol 33(8):582–594.

27. Dediu D, Levinson SC (2018) Neanderthal language revisited: not only us. Evol Lang 21:49–55.

28. Dediu D, Levinson SC (2013) On the antiquity of language: the reinterpretation of Neandertal linguistic capacities and its consequences. Front Psychol 4(397):1–17.

29. Martins PT, Marí M, Boeckx C (2018) SRGAP2 and the gradual evolution of the modern human language faculty. J Lang Evol 3(1):67–78.

30. Pääbo S (2014) The human condition—a molecular approach. Cell 157(1):216–226.

31. Kuhlwilm M, Boeckx C (2018) Genetic differences between humans and other hominins contribute to the”human condition.”

32. Reich D (2018) Who We Are and How We Got Here: Ancient DNA and the new science of the human past (Oxford University Press, Oxford).

33. de Boer B, Thompson B (2018) Biology-culture co-evolution in finite populations. Sci Rep 8(1):1209.

34. Thompson B, Kirby S, Smith K (2016) Culture shapes the evolution of cognition. Proc Natl Acad Sci 113(16):4530–4535.

35. Thomas J, Kirby S (2018) Self domestication and the evolution of language. Biol Philos 33(1-2):9.

36. Theofanopoulou C, et al. (2017) Self-domestication in Homo sapiens: Insights from comparative genomics. PloS One 12(10):e0185306.

