## Supplementary Sections for "Evolutionary Dynamics Do Not Motivate a Single-Mutant Theory of Human Language"

Bart de Boer[1]

Bill Thompson[2]

Andrea Ravignani[1,3]

Cedric Boeckx[4,5,6]

**Affiliations:**

1. AI-lab, Vrije Universiteit Brussel, Pleinlaan 2, 1050 Brussel, Belgium
2. Language and Cognition Department, Max Planck Institut für Psycholinguistik, Wundtlaan 1, 6525 XD Nijmegen, the Netherlands
3. Research Department, Sealcentre Pieterburen, Hoofdstraat 94a, 9968 AG Pieterburen, The Netherlands
4. ICREA, Passeig Lluís Companys 23, 08010 Barcelona, Spain
5. Institute for Complex Systems, Universitat de Barcelona, 585 Gran Via, 08007 Barcelona, Spain
6. Section of General Linguistics, Universitat de Barcelona, 585 Gran Via, 08007 Barcelona, Spain

**Corresponding Author:**

Bart de Boer

AI-lab, Vrije Universiteit Brussel

Pleinlaan 2

1050 Brussel

Belgium

+32 2 629 3755

**This PDF file includes:**

Supplementary text
References for SI citations

**S1. Mean and Variance Approximations**

In this section we provide three ways of formulating mean and variance in the backwards Kolmogorov equation and compare them with a direct Monte-Carlo simulation of the selection process.

The first models the population as if it were haploid. The mean change and its variance are then:

$$M_{\delta p}(p) = p(1-p)\frac{s_{aA}}{1+s_{aA}\cdot p} \tag{S.1}$$

$$V_{\delta p}(p) = \frac{p(1-p)}{2N}\frac{2+s_{aA}}{1+s_{aA}\cdot p} \tag{S.2}$$

where $s_{aA}$ is the selection coefficient of the heterozygote mutant (i.e. assuming the fitness of the wild type is 1, the fitness of the heterozygote mutant is $1+s_{aA}$) and $N$ is the population size.

A more precise approximation also considers the influence of the homozygote mutants. This leads to the following expressions:

$$M_{\delta p}(p) = p(1-p)\frac{s_{aA}+(s_{AA}-2s_{aA})p}{\bar{w}} \tag{S.3}$$

$$V_{\delta p}(p) = \frac{p(1-p)}{2N}\cdot\frac{(1+s_{aA}+(s_{AA}-s_{aA})p)(1+s_{aA}\cdot p)}{\bar{w}^2} \tag{S.4}$$

Where:

$$\bar{w} = 1+2s_{aA}p(1-p)+s_{AA}p^2 \tag{S.5}$$

is the mean fitness of the population, and $s_{AA}$ is the selection coefficient of heterozygote mutants.

A third model is due to Kimura (1980) in which it is assumed that the selection coefficients are small, such that we can ignore the effect of the mean fitness. Also, Kimura assumes the use of the effective population size, such that the following expressions are obtained:

$$M_{\delta p}(p) = p(1-p)(s_{aA}+(s_{AA}-2s_{aA})p) \tag{S.6}$$

$$V_{\delta p}(p) = \frac{p(1-p)}{2N_e} \tag{S.7}$$

where $N_e$ is the *effective population size*, i.e. the size of a randomly mating population that behaves the same as the actual population (which may not have random mating).

**Table 1: Comparison of fixation probabilities of the different models with a direct simulation for different values of N (rows) and the selection coefficient (columns). For the simulation, the 95% confidence interval is given in brackets.**

| | | 0.1 | 0.3 | 1.0 |
|---|---|---|---|---|
| **100** | *Precise* | 0.156 | 0.363 | 0.632 |
| | *Kimura* | 0.171 | 0.443 | 0.862 |
| | *Haploid* | 0.173 | 0.407 | 0.736 |
| | *MC-simulation* | 0.171 (0.163, 0.178) | 0.413 (0.404, 0.423) | 0.794 (0.786,0.801) |
| **300** | *Precise* | 0.162 | 0.368 | 0.632 |
| | *Kimura* | 0.178 | 0.448 | 0.863 |
| | *Haploid* | 0.173 | 0.407 | 0.736 |
| | *MC-simulation* | 0.173 (0.166, 0.180) | 0.417 (0.409, 0.427) | 0.781 (0.774, 0.790) |
| **1000** | *Precise* | 0.164 | 0.368 | 0.631 |
| | *Kimura* | 0.179 | 0.449 | 0.864 |
| | *Haploid* | 0.173 | 0.407 | 0.736 |
| | *MC-simulation* | 0.179 (0.171, 0.187) | 0.419 (0.411, 0.428) | 0.797 (0.788, 0.805) |

### Quality of the Approximations

In order to compare the quality of the different approximations, they were compared to a direct simulation of the population. This simulation modeled a population that is repeatedly replaced by a new population in which the new individuals descend from two individuals in the previous population, each of which was selected with a probability that was proportional to their fitness. The fitness was 1 for the wild type, $1+s_{aA}$ for the heterozygote mutant and $1+s_{AA}$ for the homozygote mutant.

The fixation probabilities are presented in table 1. It shows them for three different population sizes (100, 300 and 1000) and three different selection coefficients (0.1, 0.3 and 1.0) that were equal for homozygotes and heterozygotes ($s_{aA} = s_{AA}$) – as was pointed out in section 2.1, this corresponds to the mutation being dominant, and this appears the most likely interpretation for Berwick and Chomsky's single-mutant hypothesis. It is clear from these results that none of the models is entirely accurate, that accuracy increases with decreasing selection coefficient and increasing population (as expected), and that the simplest approximation (assuming a haploid population) works surprisingly well. Larger population sizes were not simulated, because they were impractical, and because they were not expected to change the conclusions (accuracy increases with population size, so differences between models would become less pronounced). Smaller selection coefficients were also not simulated, because all models have good accuracy for small selection coefficients. As was pointed out in section 2.1, higher selection coefficients were considered extremely unlikely.

**S2. Diffusion Approximation of the Moran Process**

The equations that are used for investigating fixation probabilities for the haploid approximation (equations 1.4 and 1.5) in the main text are derived from the Moran process. The Moran process (Moran, 1958; Nowak, 2006, ch. 6) is a birth-death process in a finite population of size $M$ where at each time step, one member of the population dies, and one new member of the population gets born. The population consists of residents and mutants, and its state can be fully described by a single integer, the number of mutants: $0 \leq i \leq N$. At each birth death event, the number of mutants may stay the same, increase by one or decrease by one. If residents have fitness 1 and mutants have fitness $1+s_{aA}$, then the probabilities of these transitions are as follows:

$$p\left(i+1\middle|i\right)=\frac{M-i}{M}\cdot\frac{i\left(1+s_{aA}\right)}{M-i+i\left(1+s_{aA}\right)}=\frac{M-i}{M}\cdot\frac{i}{M}\cdot\frac{1+s_{aA}}{1+\frac{i}{M}s_{aA}} \tag{S.8}$$

and:

$$p\left(i-1\middle|i\right)=\frac{i}{M}\cdot\frac{M-i}{M-i+i\left(1+s_{aA}\right)}=\frac{i}{M}\cdot\frac{M-i}{M}\cdot\frac{1}{1+\frac{i}{M}s_{aA}} \tag{S.9}$$

where in both cases the first formulation illustrates that the probability is just the product of one type dying and the other being born (taking into account the fitness) and the second formulation is a more convenient form for further analysis.

By making the substitution $p\equiv i/M$ (*i.e.* $p$ represents the proportion of mutants), the equations can be rewritten to:

$$p\left(p+\frac{1}{M}\middle|p\right)=p\left(1-p\right)\cdot\frac{1+s_{aA}}{1+p\cdot s_{aA}} \tag{S.10}$$

$$p\left(p-\frac{1}{M}\middle|p\right)=p\left(1-p\right)\cdot\frac{1}{1+p\cdot s_{aA}} \tag{S.11}$$

These can be incorporated in the mean and variance terms of the backward Kolmogorov equation. By definition, they are as follows:

$$M_{\delta p}\left(p\right)=\int\delta p\cdot w\left(p+\delta p\middle|p\right)d\left(\delta p\right) \tag{S.12}$$

$$V_{\delta p}\left(p\right)=\int\left(\delta p\right)^2\cdot w\left(p+\delta p\middle|p\right)d\left(\delta p\right) \tag{S.13}$$

where $w\left(p+\delta p\middle|p\right)$ is the probability per unit time to go from a state with p mutants to one with $p+\delta p$ mutants. For a one-step process (i.e. a process where composition of the underlying discrete population changes by maximally one at each time step) like the Moran process, $w$ can be formulated as follows:

$$w\left(q\middle|p\right)=\alpha\left[\delta\left(q-p+\frac{1}{M}\right)p\left(p-\frac{1}{M}\middle|p\right)+\delta\left(q-p-\frac{1}{M}\right)p\left(p+\frac{1}{M}\middle|p\right)\right] \tag{S.14}$$

where $\alpha$ is a conversion constant that converts the continuous time variable of the diffusion process to the discrete time variable of the Moran process, and $\delta(\ x\ )$ is the Dirac delta function. Substituting this in equations (S.12) and (S.13) gives:

$$M_{\delta p}=\frac{\alpha}{M}\left[p\left(p+\frac{1}{M}\middle|p\right)-p\left(p-\frac{1}{M}\middle|p\right)\right] \tag{S.15}$$

$$V_{\delta p}=\frac{\alpha}{M^2}\left[p\left(p+\frac{1}{M}\middle|p\right)+p\left(p-\frac{1}{M}\middle|p\right)\right] \tag{S.16}$$

These formulations are still general for any one-step process. Substituting (S.10) and (S.11) gives the formulation for the Moran process:

$$M_{\delta p}=\frac{\alpha}{M}p\left(1-p\right)\frac{s_{aA}}{1+p\cdot s_{aA}} \tag{S.17}$$

$$V_{\delta p}=\frac{\alpha}{M^2}p\left(1-p\right)\frac{2+s_{aA}}{1+p\cdot s_{aA}} \tag{S.18}$$

Choosing $\alpha = M$ ensures that time is expressed in generations (i.e. the number of time steps needed to replace a number of agents that is equal to the population size). Finally, because we are modeling a diploid population (admittedly pretending we can do that by only looking at the heterozygotes) we need to choose $M = 2N$, and we obtain equations 1.4 and 1.5 of the main text.

### S3. Sensitivity of number of mutations to parameters *m* and α

The relationship between the slope of the exponential distribution that describes the probability of mutations that provide a given positive selection coefficient ($-\alpha$), the total improvement that needs to be achieved ($I$) and the expected number of mutations (‖m‖) has been calculated through drawing random samples from the appropriate distribution (equation 15 in the main text) for different values of $\alpha$ and $I$. The mean value of the needed improvements is given in figure S3.1.

For larger numbers of mutations, this number can be approximated with satisfactory accuracy by dividing the total improvement by the mean size of a mutation that goes to fixation. This is given by:

$$\left\|m\right\| \approx I \cdot \frac{\alpha\left(2+\alpha\right)}{2+2\alpha} \qquad \text{(S.19)}$$

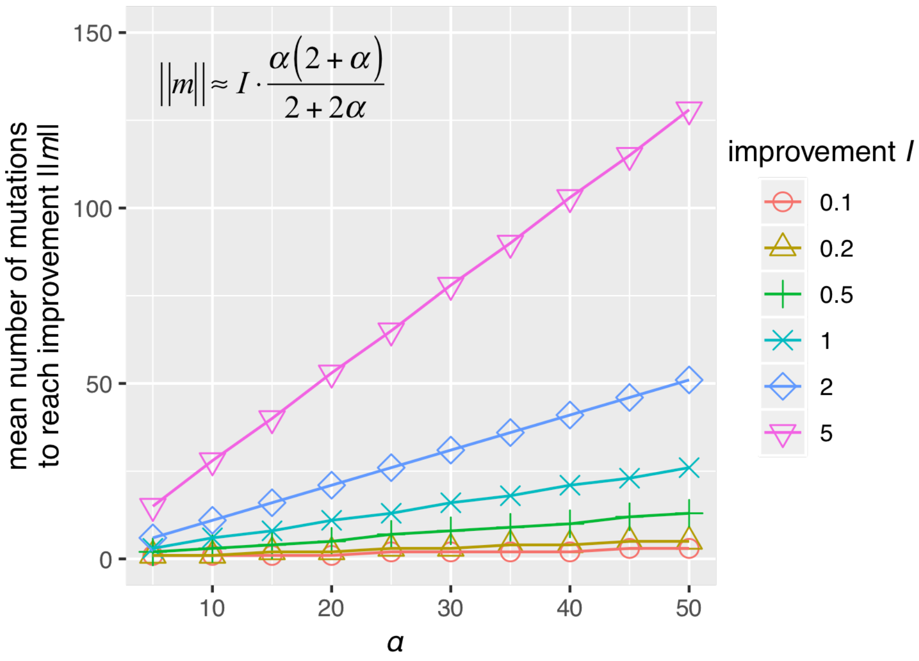


**Figure S3.1: Mean number of improvements needed to reach a given size of improvement *I* for different values of the slope the distribution of mutation sizes –*α*. The approximate equation for this relation is given in the top left corner of the graph.**

### References

Moran, P.A.P. (1958). Random processes in genetics. *Mathematical Proceedings of the Cambridge Philosophical Society*, 54(1), 60-71.

Nowak, M.A. (2006). *Evolutionary dynamics*. Cambridge, MA: Harvard University Press.
